# Efficient hydrogen-dependent carbon dioxide reduction by *Escherichia coli*

**DOI:** 10.1101/169854

**Authors:** Magali Roger, Fraser Brown, William Gabrielli, Frank Sargent

**Affiliations:** School of Life Sciences, University of Dundee, DUNDEE DD1 5EH, Scotland; Ingenza Ltd, Roslin Biocentre, EDINBURGH EH25 9PP, Scotland; Sasol UK Ltd, St Andrews Laboratory, North Haugh, ST ANDREWS KY16 9ST, Scotland

**Author notes:** For correspondence: Prof Frank Sargent FRSE, Division of Molecular Microbiology, School of Life Sciences, University of Dundee, Dow Street, DUNDEE DD1 5EH, Scotland.

## Abstract

The formate hydrogenlyase (FHL) complex of *Escherichia coli* is normally produced under anaerobic fermentative conditions when its physiological role is to oxidise formate to carbon dioxide and couple that reaction directly to the reduction of protons to molecular hydrogen. This forward reaction is the major route of hydrogen production by *E. coli* and the physiology, genetics and biochemistry of this process are reasonably well understood. However, studies of the reverse reaction - hydrogen-dependent carbon dioxide reduction – have been rather less extensive. Harnessing the alternative reverse reaction has the potential to unlock FHL as a carbon dioxide cycling enzyme, or could potentially lead to the development of bio-based carbon capture technologies. In this work, it is established that FHL can operate as a highly efficient CO_2_ reductase. A controllable pressure system was designed with the intention of maximising substrate availability to the FHL enzyme. By placing gaseous CO_2_ and H_2_ are under pressure (up to 10 bar), and by using a compartmentalised intact whole cell approach where the produced formate is excreted, the optimised experimental system was observed to convert 100 % of gaseous CO_2_ to formic acid and generate >500 mM formate in solution at an initial rate of 1.2 g formate *per* litre *per* hour.

## INTRODUCTION

Hydrogen-dependent reduction of carbon dioxide (CO_2_) to formate (HCOO^−^) offers a promising route to greenhouse gas sequestration as well as development of carbon abatement technologies, hydrogen transport and storage, and the sustainable generation of renewable chemical feedstocks (Kamm *et al*., 2006). The most common approach to performing direct hydrogenation of CO_2_ to formate is to use chemical catalysts in homogeneous or heterogeneous reactions requiring catalysts together with extreme reaction conditions (Appel *et al*., 2013, Aresta & Dibenedetto, 2007). An alternative approach is to use the ability of living organisms to perform this reaction biologically. However, although CO_2_ fixation pathways are widely distributed in nature, few enzymes have been described that have the ability to perform the direct hydrogen-dependent reduction of CO_2_ (Schuchmann & Muller, 2013, Alissandratos *et al*., 2014).

Under anaerobic conditions *Escherichia coli* can carry out a mixed-acid fermentation of glucose to produce ethanol and a variety of organic acids, one of which is the one-carbon compound formate. The formate so-produced normally accumulates outside the cell until its rising concentration begins to cause a drop in extracellular pH. When the pH reaches <6.8 the formate is transported back into the cell and induces assembly of the seven subunit, membrane-bound formate hydrogenlyase (FHL) complex (Figure 1a) (Sargent, 2016). Under these conditions the thermodynamics are favourable for hydrogen gas production (the ‘forward reaction’), with the predicted small Gibb’s free energy release, together with volatile (H_2_) or rapidly transformable (CO_2_) reaction products, allowing rapid detoxification of the organic acid (McDowall *et al*., 2014).

**Figure 1:**
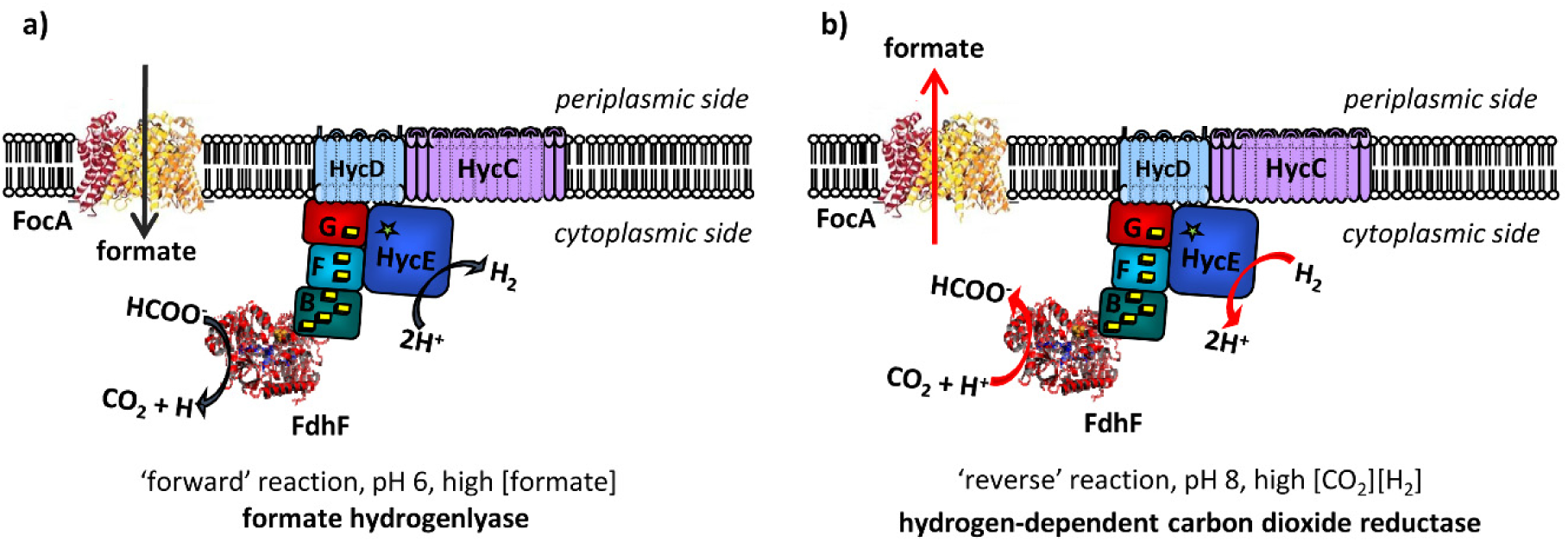
The two possible reactions performed by the *E. coli* FHL complex. **(a)** Under physiological conditions, formate produced anaerobically is initially excreted. When extracellular levels increase and the pH drops below 7, formate is taken up through the FocA channel and induces synthesis of the FHL complex, which oxidises the formate and produces H_2_ gas. **(b)** Cells already containing the FHL complex can efficiently perform a hydrogen-dependent CO_2_ reductase reaction when placed under permissive conditions.

*E. coli* FHL comprises a molybdenum-dependent formate dehydrogenase linked by Fe-S cluster-containing-proteins to a [NiFe]-hydrogenase, termed Hyd-3 (Figure 1a). This catalytic domain is anchored at the cytoplasmic side of the membrane by the HycCD membrane proteins (Figure 1a). The requirement for membrane attachment of the catalytic domain is not fully understood, however the enzyme shares an evolutionary lineage with those involved in generating transmembrane ion gradients (Nitschke & Russell, 2009, Batista *et al*., 2013, Lim *et al*., 2014). There is, however, no direct evidence that *E. coli* FHL itself couples formate oxidation to ion transport (Pinske & Sargent, 2016).

FHL is never normally synthesised by *E. coli* in the absence of formate, so its ability to perform the ‘reverse reaction’ (Figure 1b) has not been extensively explored. However, other bacteria, such as the Gram-positive spore-forming acetogen *Acetobacterium woodii*, have been shown to grow under a H_2_ / CO_2_ atmosphere and a hydrogen-dependent CO_2_ reductase (HDCR) was recently identified in that organism (Schuchmann & Muller, 2013). The *A. woodii* HDCR was found to be a molybdenum-dependent formate dehydrogenase linked to an electron bifurcating [FeFe]-hydrogenase (*i.e.* a hydrogenase from a different family to *E. coli* Hyd-3). In the presence of uncouplers, the *A. woodii* HDCR allowed the cells to generate formate from gaseous substrates (Schuchmann & Muller, 2013). Very early work suggested that a version of *E. coli* could potentially generate some formate from H_2_ and CO_2_ (Woods, 1936), although of course this research was carried out in an era before modern genetic techniques could verify that FHL was directly responsible, and more recently the purified formate dehydrogenase component of *E. coli* FHL was reported to be reversible when attached to an electrode (Bassegoda *et al*., 2014). Initial experiments with intact *E. coli* cells first suggested the Hyd-3 component of FHL was reversible with artificial electron carriers (Sawers *et al*., 1985, Maeda *et al*., 2007), before electrochemical studies of purified *E. coli* Hyd-3 established a *K*_m_ for H_2_ of 34 μM (McDowall et al., 2014). Subsequently, intact *E. coli* cells pre-grown to induce FHL and then washed and incubated with H_2_ and CO_2_ under atmospheric pressure, were able to generate low levels of formate in an FHL-dependent manner (Pinske & Sargent, 2016), so proving the principle that *E. coli* FHL could act as a hydrogen-dependent CO_2_ reductase when operating in ‘reverse’ (Figure 1b). However, the efficiency of the reaction under the original test conditions remained sub-optimal (maximum 18 % of gaseous CO_2_ converted to formate) (Pinske & Sargent, 2016).

In this work, experiments with pressurised gases reveal the optimum reaction conditions that allow efficient hydrogen-dependent CO_2_ reduction be *E. coli.* By placing a H_2_:CO_2_ mixture under a constant 10 bar pressure above an *E. coli* cell suspension, it is demonstrated here that >0.5 M formate can be generated in solution representing 100 % conversion of gaseous carbon dioxide.

## RESULTS

### Modelling gas solubility in aqueous solution

It is thought that CO_2_ itself, as opposed to carbonate (CO_3^2-^_) or bicarbonate (HCO_3^−^_), is the direct product (and substrate) for bacterial formate dehydrogenase (Fdh) enzymes (Thauer *et al*., 1977). At neutral pH, the solubility of CO_2_ is known to be low and its behaviour in solution is known to be complex (Carroll *et al*., 1991), thus substrate availability to the FHL enzyme is likely to be a limiting parameter. As it has been suggested for some industrial bioprocessing studies, the used of increased headspace gas pressure could be an approach to enhancing gas solubility in aqueous solution since Henry’s Law states that the amount of dissolved gas should be proportional to the applied pressure Lopes *et al*., 2014). To begin to test this hypothesis, it was first investigated what relative concentrations of dissolved H_2_ and CO_2_ might be attainable by applying headspace pressure to a 1:1 mixture of these gases. To do this, a non-random two-liquid (NRTL) activity coefficient model with Henry’s law for H_2_ and CO_2_ derived from isothermal data sets at 308 K/35 C was devised (Figure 2). The model was sufficient to predict H_2_ and CO_2_ solubilities in water at low concentrations, or in this case for pressures below 20 bar. At isothermal conditions (308 K/35 C) at neutral pH, the correlation of solubility (mol/m^3^ ≡ mmol.L^−1^) and absolute pressure (bar) follows y = 12.61x and y = 0.3651x for CO_2_ and H_2_, respectively (Figure 2). The model, consistent with Henry’s Law, predicts that an almost 10-times increase in solubility can be attained at 10 bar *versus* 1 bar for both H_2_ and CO_2_. In terms of absolute values, the model predicts CO_2_ could reach ~120 mmol.L^−1^ in solution, and H_2_ ~4 mmol.L^−1^, when mixed together at 10 bar pressure (Figure 2).

**Figure 2:**
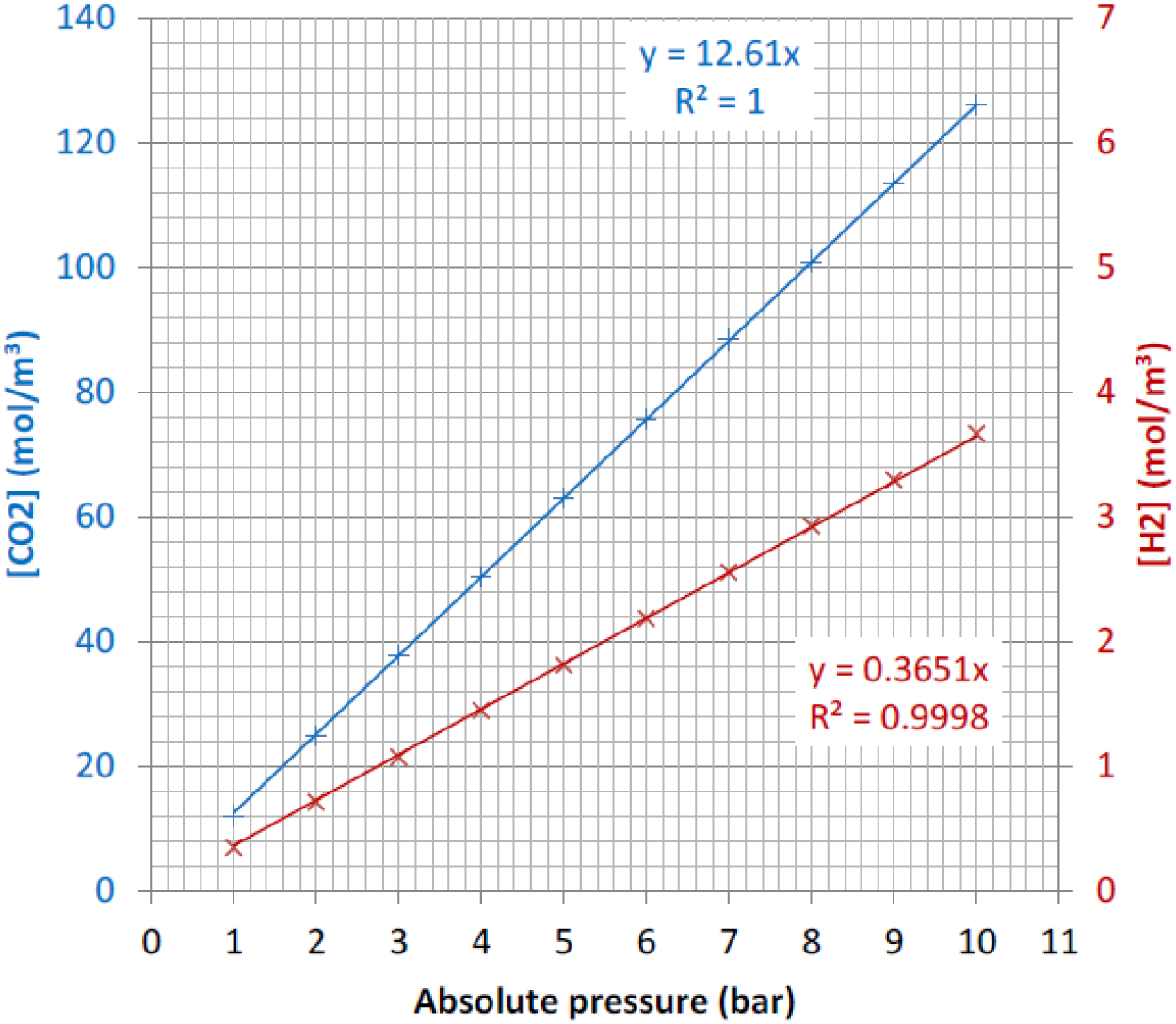
The relationship between CO_2_ and H_2_ solubilities in water for pressures up to 10 bar. The non-random two-liquid (NRTL) activity coefficient model with Henry’s law for H_2_ and CO_2_ was derived from published isothermal data sets for H_2_ and CO_2_ at 35 ^<u>°</u>^C using Aspen Plus. The model assumes a 1:1 mixture of CO_2_:H_2_ under the given conditions.

### Designing a pressurised bioreactor for CO_2_ uptake/formate production experiments

Next, in order to test experimentally the effect of utilising pressurised gases as substrates, a high-pressure bioreactor system was designed with the aim of catalysing the direct hydrogen-dependent reduction of CO_2_ to formate using intact bacterial cells (Figure 3). This bioreactor system was composed of two stirred steel reactors (Figure 3). The pre-mixing “H_2_:CO_2_ ballast vessel” allowed the preparation of a homogeneous gas mixture (~44 % H_2_ and ~56 % CO_2_ as quantified by Gas Chromatography) at high pressure (40 bar). This vessel was then used for the pressurisation of the “production vessel”, which was the bioreactor containing the bacterial cell suspension that initiated the biological reaction (Figure 3). The system was designed with the ability to operate at constant temperatures, to monitor and modify the pH in the production vessel, to monitor gas consumption in the ballast vessel, and to withdraw liquid samples from the production vessel for analysis.

**Figure 3:**
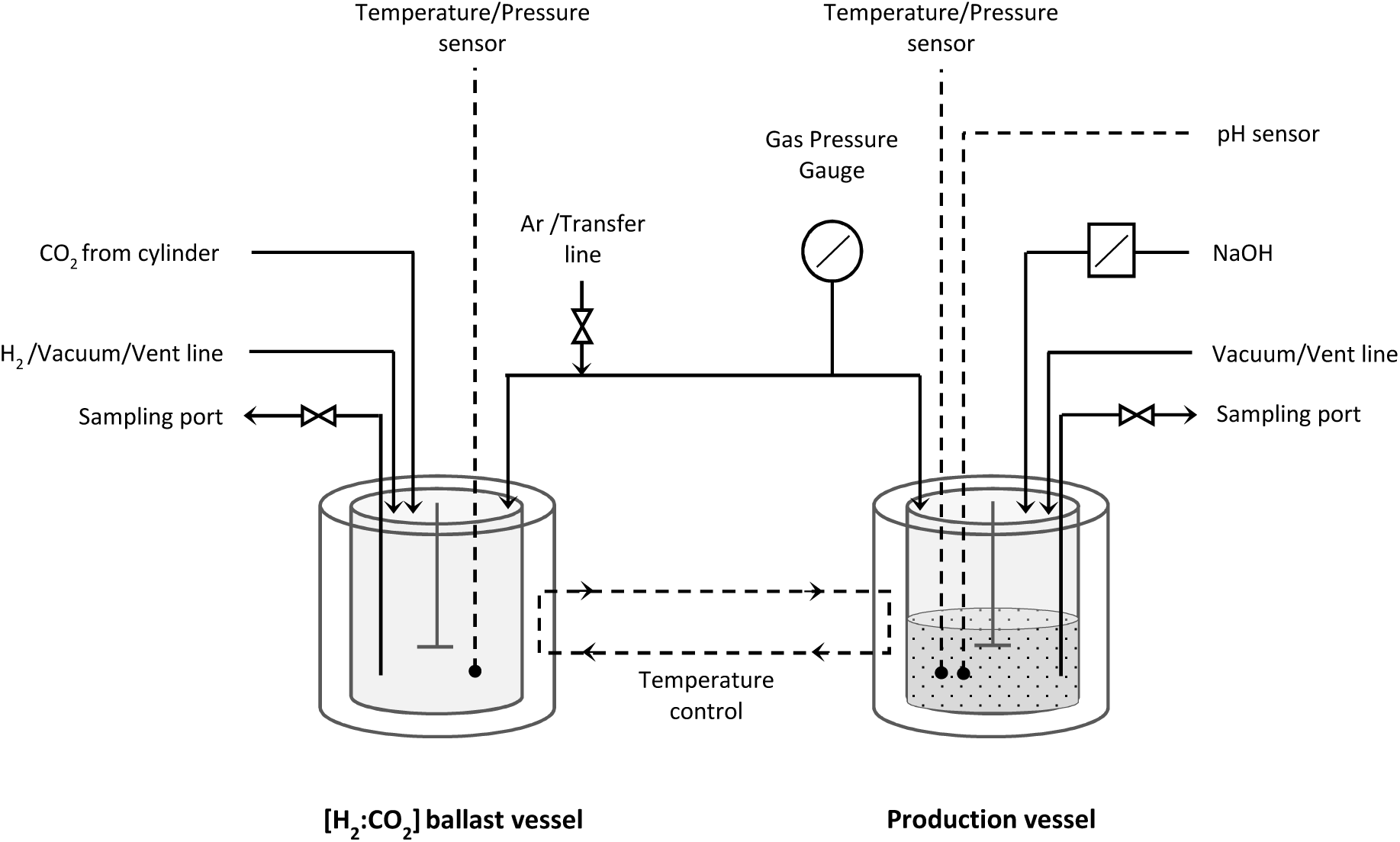
Experimental setup for the bioconversion of CO_2_ and H_2_ into formate using high-pressure reactors. The stainless steel ballast vessel is used for pre-pressurisation of gas mixtures (at 40 bar) and a sampling port allows the amount of each gas in the mixture to be accurately recorded. The ballast vessel is connected to a stainless steel production vessel. This contains the cell suspension in buffer that can be placed under constant pressure from the ballast vessel. It is possible to monitor and control pH in the production vessel and sample the aqueous phase.

### Application of gas pressure increases the efficiency of the FHL reverse reaction

The *E. coli* strain FTD89, which has a genotype of Δ*hyaB/*Δ*hybC* and thus lacks all major hydrogenase activity except that from FHL, was grown under anaerobic fermentative conditions in order to induce synthesis of the FHL complex. The intact whole cells were then harvested and washed extensively before being placed in a solution containing only 20 mmol.L^−1^ MOPS buffer (pH 7.4) at 25 g wet weight cells.L^−1^. This cell suspension was then placed in the production vessel (Figure 3) under a constant 2 bar pressure of H_2_:CO_2_ mixture (44:56 ratio as calculated by gas chromatography), corresponding to a constant 27.52 mmol.L^−1^ CO_2_ and 0.81 mmol.L^−1^ H_2_ in the aqueous phase. The increase in concentration of formate was then followed over time by High Performance Liquid Chromatography (HPLC) (Figure 4 **and** Figure 5a), while the decrease in ballast vessel gas pressure, indicating gas consumption in the production vessel (Figure 5b) and the pH changes in the production vessel (Figure 5c) were all similarly monitored. Under these 2 bar/MOPS pH 7.4 conditions the concentration of formate in the cell suspension was observed to initially increase and then level off after a few hours with a final concentration of formate produced in the reaction vessel of 8 mmol.L^−1^ (Figure 5a). However, this was concomitant with a strong decrease in the pH in the production vessel (Figure 5c), which can be attributed to both CO_2_ dissolution (at the beginning of the experiment) as well as production of formate in the aqueous phase.

**Figure 4:**
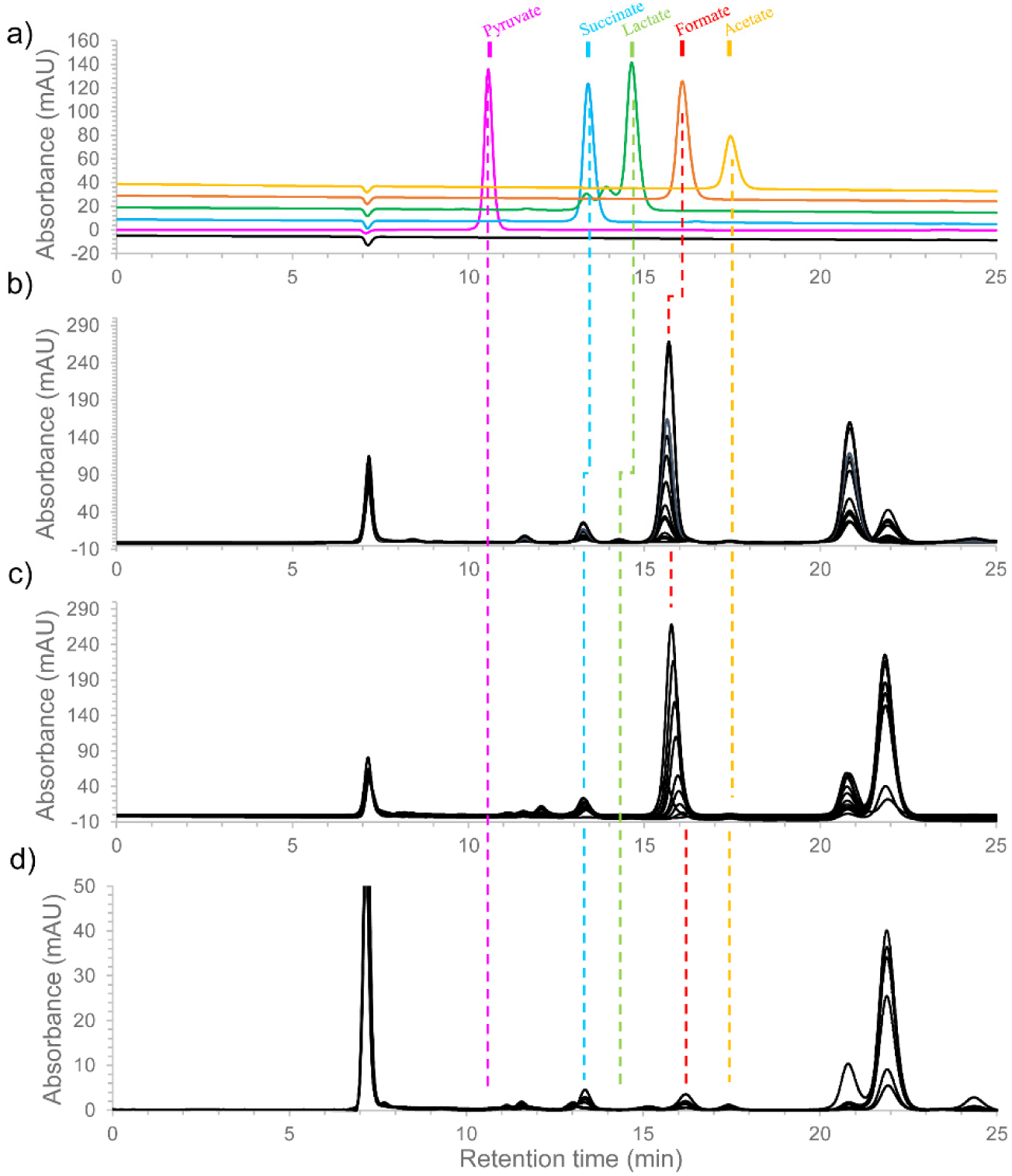
HPLC detection of organic acids produced in the cell suspension during hydrogen-dependent CO_2_ reduction. **(a)** 10 μL of various organic acids standards or clarified samples taken from the production vessel containing either *E. coli* FTD89 **(b)**, RT1 **(c)** or the control FHL-minus strain RT2 **(d)** at different time points of the reaction were applied to an Aminex HPX 87H column at 50^<u>°</u>^C, using sulfuric acid as mobile phase (0.5 mL.min^−1^). Separated compounds were detected at A_210nm_. A trace of succinate (13.4 min retention time) and formate (16.2 min retention time) can be detected at the beginning of the experiment. Following pressurisation of the production vessel containing FTD89 or RT1 washed whole-cells with a constant ratio of H_2_:CO_2_ at 2 bar pressure, formate increased over the time, while the RT2 strain was unable to generate formate from H_2_ and CO_2_. Note that two peaks can be observed around 21 and 22 min retention time. Both of them have already been observed in previous study (Pinske & Sargent, 2016), however attempts to identify these two peaks gave no clear results and their identities remain to be determined. The two peaks were also observed after incubation of FHL-minus control strain RT2 **(d)** demonstrating that these compounds are produced with FHL-independent manner.

**Figure 5:**
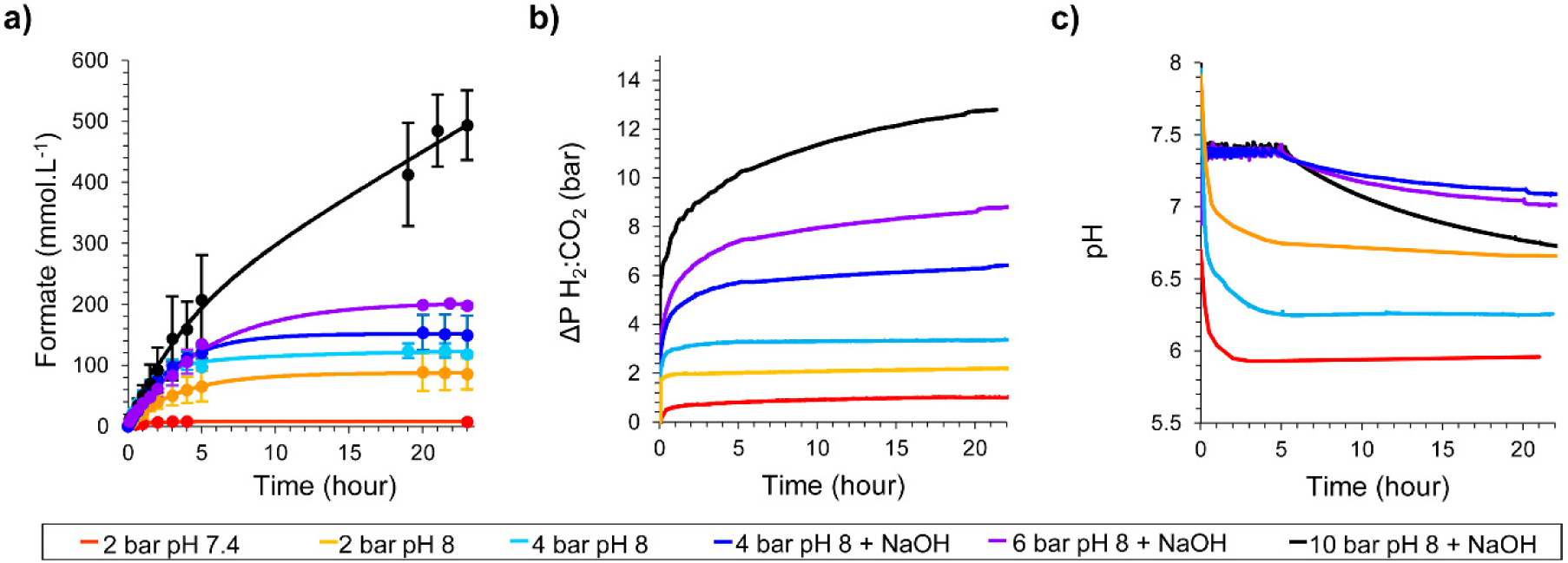
Hydrogen-dependent CO_2_ reduction to formate using a high-pressure reactor. Cultures of *E. coli* FTD89 strain (Δ*hyaB*, Δ*hybC*) were pre-grown under FHL-inducing conditions before 25 g of washed, intact whole-cells were placed in a high pressure reactor and incubated at a constant H_2_:CO_2_ ratio (~1:1) at 2, 4, 6 or 10 bar pressure in a final volume of 500 mL at 37 °C with stirring at 500 rpm. **(a)** Formate production in the production vessel was recorded over time by manual sampling and quantification by HPLC. **(b)** The pressure decrease in the gas mixing ballast vessel was recorded over time under different pressure conditions applied to the production vessel. **(c)** The pH in the production vessel was monitored over the time course of the reaction under different gas pressures. The colour key informs on the different gas pressures and buffering conditions applied. Samples at pH 7.4 were in 20 mM MOPS buffer; samples at pH 8 were in 200 mM Tris.HCl buffer; and samples labelled ‘+NaOH’ were titrated with 2M NaOH during the reaction.

In order to minimise the pH drop observed upon gas pressurisation and formate production, the pH of the starting buffer was increased from pH 7.4 to pH 8.0 and MOPS buffer was replaced by 200 mmol.L^−1^ Tris.HCl. At 2 bar pressure, these modifications alone resulted in 85 mmol.L^−1^ for the final concentration of formate produced (Figure 5a), and increasing the gas pressure to 4 bar allowed a further increase of the final concentration of formate produced to 120 mmol.L^−1^ (Figure 5a).

Next, the production vessel was further modified to allow the addition of sodium hydroxide to the *E. coli* cell suspension in order to maintain the pH above 6.8 during the reaction. By using this strategy, a further increase in the final formate concentration to 150 and 200 mmol.L^−1^ was observed at 4 and 6 bar pressure, respectively (Figure 5a). Finally, increasing the pressure to 10 bar, which would result in 122.88 mmol.L^−1^ CO_2_ and 3.61 mmol.L^−1^ H_2_ in solution, together with the continuous pH regulation system in operation, allowed the production of >0.5 mol.L^−1^ formate in the bioreactor over the 23 hours’ time-course of the experiment (Figure 5a).

It can be concluded from these experiments that maintaining the pressure of gas mixture in the headspace at 10 bar, combined with the fine control of the reaction pH, leads to an over 20-times increase in the total amount of formate produced *per* mg of total cell protein *versus* that observed at ambient pressure (Figure 5a). Indeed, the efficiency of this reaction was observed to be optimal, with a value of 103.0 % conversion of gaseous CO_2_ to formate in solution recorded at 10 bar pressure (Figure 6). The reaction is dependent upon the presence of the FHL complex in the cells, with a mutant strain (RT2) devoid of the genes encoding the enzyme being unable to generate formate (Figure 4d). Intact *E. coli* cells are therefore, under the correct conditions, capable of a highly efficient hydrogen-dependent reduction of CO_2_ to formate.

**Figure 6:**
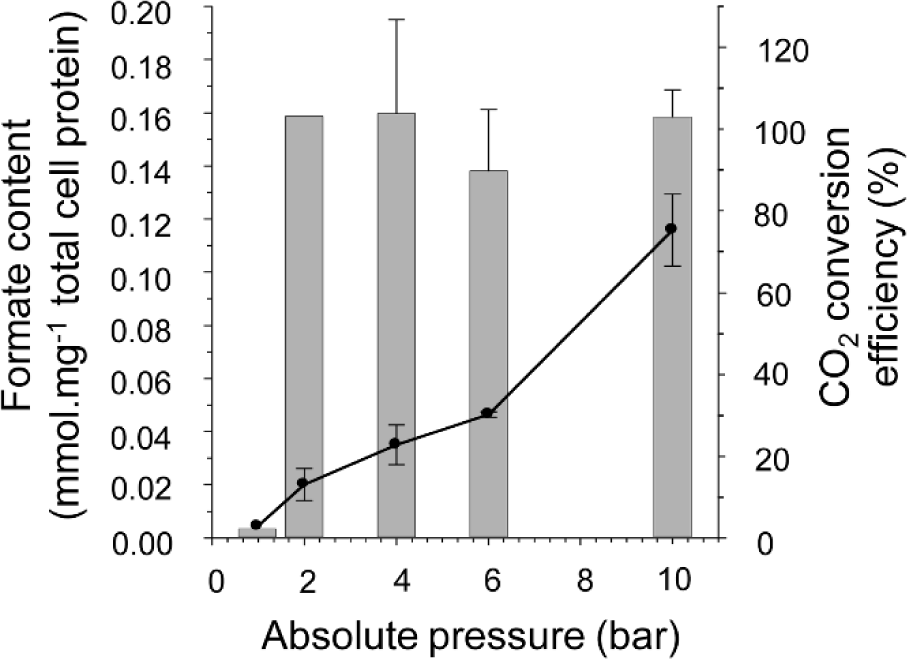
Final concentrations of formate produced under pressure. A comparison of the final formate content of the production vessel under different gas pressures (left x-axis, dotted line) with the overall efficiency of CO_2_ conversion to formate by calculating and comparing CO_2_ uptake and formate production levels (right x-axis, grey bars).

### Towards strain optimisation for hydrogen-dependent CO_2_ reduction

The *E. coli* FTD89 strain utilised thus far contains, in addition to FHL, two other Fdhs (Sawers, 1994) and the potential ability to assimilate some of the formate produced through the reverse reaction of pyruvate formatelyase (PFL) (Zelcbuch *et al*., 2016). Although in the current reaction conditions there are no exogenous respiratory electron acceptors, so the other Fdhs should not be functional, and there are no exogenous carbon sources, so acetyl CoA levels should be minimal, it was considered that genetic inactivation of potential fomate utilisation pathways may help optimise the CO_2_ reduction to this organic acid. Therefore, the ability of an additionally-modified *E. coli* strain RT1 (Δ*hyaB*, Δ*hybC*, Δ*pflA*, Δ*fdhE*) to perform hydrogen-dependent CO_2_ reduction was compared to FTD89. In RT1, the *fdhE* mutation inactivates biosynthesis of the respiratory Fdhs but does not affect the enzyme associated with FHL (Schlindwein *et al*., 1990, Luke *et al*., 2008) and the *pflA* mutation removes the PFL activating enzyme (Sawers & Watson, 1998).

Using low-pressure, small-scale experiments, as shown in Figure 7, a two-times increase in the final amount of formate produced from gaseous H_2_ and CO_2_ can be recorded when using a suspension of the *E. coli* strain RT1 in comparison with the FTD89 strain. An *E. coli* control strain RT2 (Δ*hyaB*, Δ*hybC*, Δ*pflA*, Δ*fdhE*, Δ*hycA*-*I*), which is genetically identical to the RT1 strain but further deleted for the *hycABCDEFGHI* operon encoding the Hyd-3 [NiFe]-hydrogenase component of FHL, could not produce formate under the same conditions (Figure 7).

**Figure 7:**
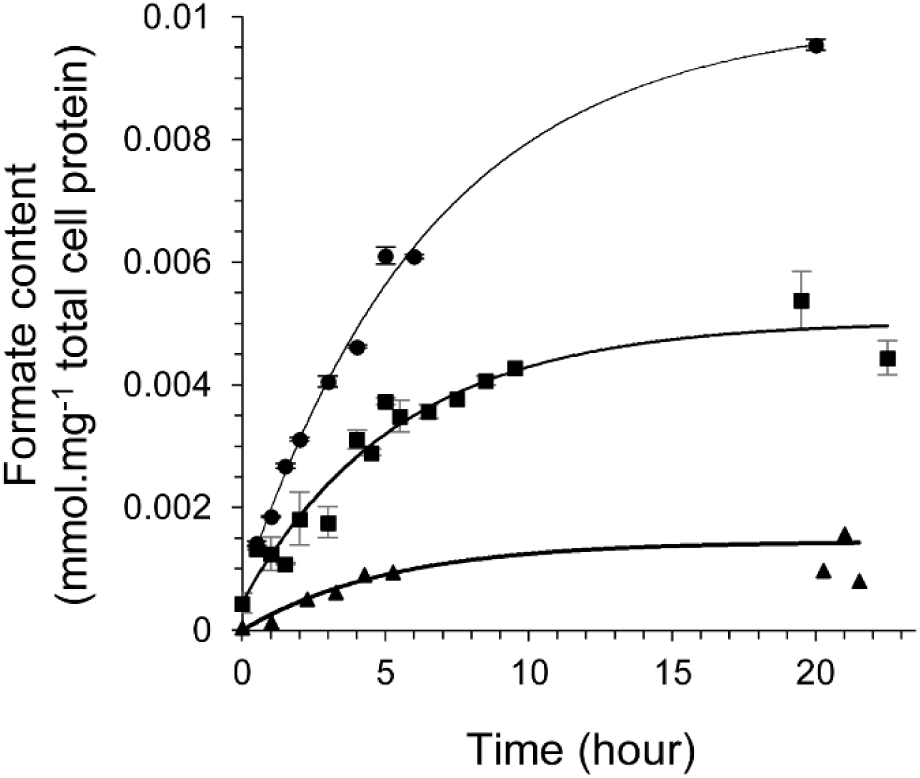
Optimising CO_2_ bioconversion by genetic engineering and low-pressure assays. Cultures of *E. coli*strains FTD89 (Δ*hyaB*, Δ*hybC*) [black square], RT1 (Δ*hyaB*, Δ*hybC*, Δ*pflA*, Δ*fdhE*) [black circle] and RT2 (Δ*hyaB*, Δ*hybC*, Δ*pflA*, Δ*fdhE*, Δ*hycA*-*I*) [black triangle] were pre-grown under FHL-inducing conditions. Then small-scale 25 mg samples of washed whole-cells were incubated in sealed Hungate tubes in a final volume of 3 mL 20 mmol.L^−1^ MOPS buffer pH 7.4 at 37 °C under a CO_2_ and H_2_ atmosphere at ambient pressure. The formate concentration in the liquid phase of the reaction tubes was assayed by HPLC over time.

### Optimising biomass requirements for hydrogen-dependent CO_2_ reduction

Attention next returned to the high pressure bioreactor and the *E. coli* RT1 strain (Δ*hyaB*, Δ*hybC*, Δ*pflA*, Δ*fdhE*) was used to further explore the optimal conditions for hydrogen-dependent CO_2_ reduction (Figure 8). To establish the optimum amount of biomass necessary for efficient hydrogen-dependent reduction of CO_2_, different amounts of intact *E. coli* RT1 cells (2, 4, 8, 16, 25 and 50 g wet weight L^−1^), pre-grown to induce FHL expression, were incubated in the 500 mL reaction vessel at a constant H_2_:CO_2_ pressure of 10 bar and the final concentrations of formate produced in the aqueous phase of the bioreactor, and its initial rate of production over time, were determined (Figure 8). When the amount of cell protein used is taken into account (Figure 8a), the greatest relative final concentration of formic acid is achieved when the RT1 cells were prepared at 8 g.L^−1^ (Figure 8a). This amount of cells also corresponded to the point where conversion of CO_2_ to formic acid reached optimum efficiency (Figure 8b). Indeed, increasing the RT1 biomass beyond 8 g.L^−1^ up to 50 g.L^−1^ (25 g cells, wet weight, in the 500 mL reaction vessel) did not contribute to an increase in the final amounts of formate produced (Figure 8a,b).

**Figure 8:**
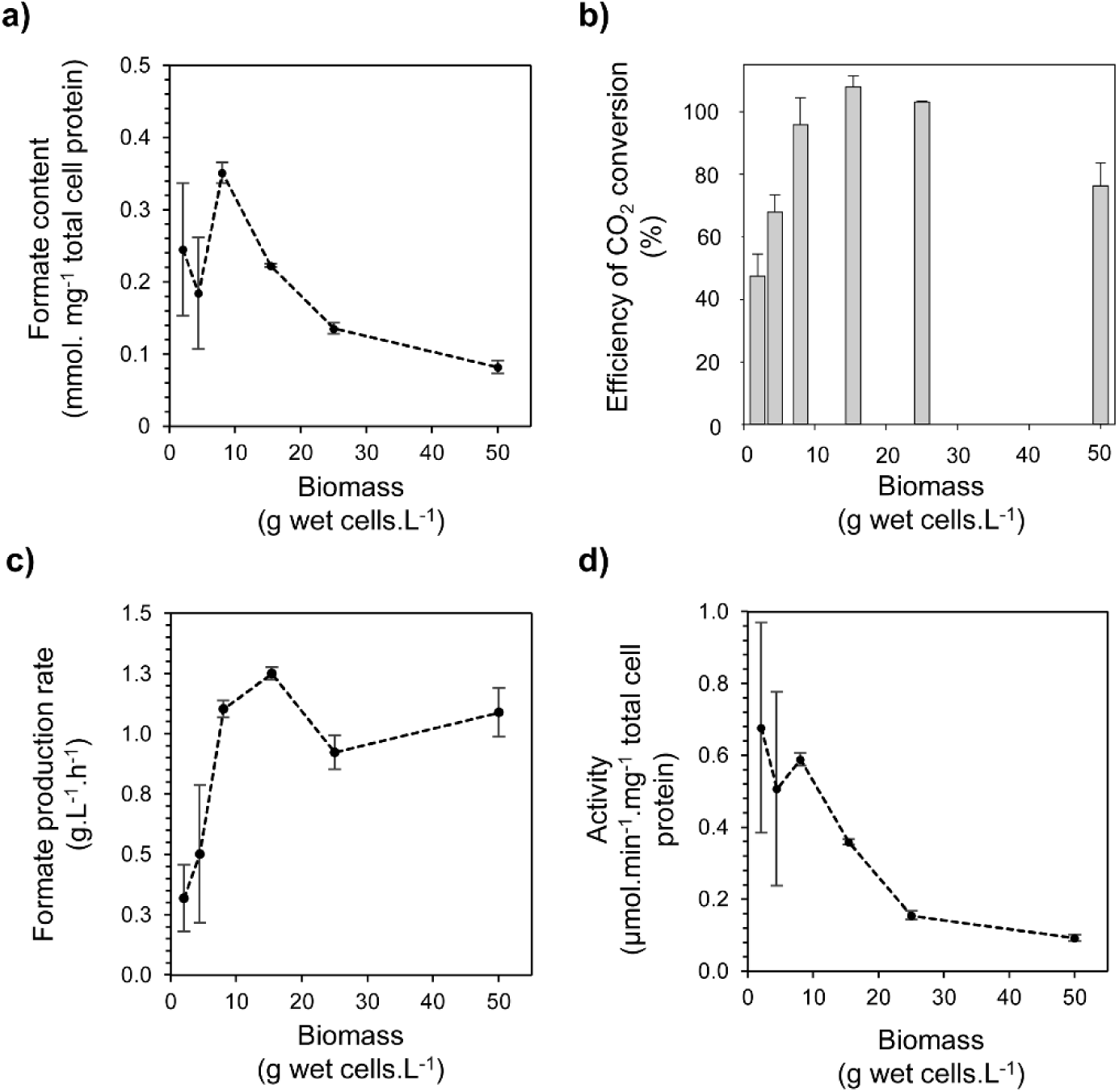
Optimising biomass requirements for formate production from CO_2_ at 10 bar. Cultures of *E. coli* RT1 strain (Δ*hyaB*, Δ*hybC*, Δ*pflA*, Δ*fdhE*) were pre-grown under FHL-inducing conditions. Various amounts (2, 4, 8, 16, 25, 50 g wet weight L^−1^) of washed whole-cells were incubated at constant H_2_:CO_2_ (~1:1) at 10 bar pressure in a final volume of 500 mL at 37 °C and 500 rpm in the high pressure reaction vessel. Formate production over the time course of the reaction was recorded by manual sampling and quantified by HPLC. **(a)** The total formate content in the production vessel at the end of the reaction (23 hours) as a factor of total cell protein used. **(b)** The apparent efficiency of CO_2_ conversion to formate as calculated by comparing CO_2_ uptake with formate production. **(c)** The initial rates of formate production under different conditions calculated by extrapolating formate production time-courses, and **(d)** overall ‘activity’ of the FHL-dependent formate production pathway by incorporating the protein concentrations present in each reaction with the initial rates calculated in panel (c).

In terms of the initial rates of formate production (Figure 8c,d), increasing the amount of RT1 cells allowed a clear increase in the apparent rate of formate production at 10 bar pressure (Figure 8c), which stabilised at ~ 1.2 g formate produced L^−1^ hr^‐1^ through 8-16 g.L^−1^ cells (Figure 8c). When these initial formate production rates are calculated by taking into account the relative protein concentrations present in the reactions (termed ‘activity’ in Figure 8d), it is also clear that 8 g. L^−1^ of RT1 cells are optimum under these conditions, with an initial rate of 0.6 μmol formate produced. min^−1^. mg^−1^ total cell protein.

## DISCUSSION

### Formate hydrogenlyase as a hydrogen-dependent CO_2_ reductase

Disproportionation of formate to CO_2_ and H_2_ by FHL (termed as the ‘forward reaction’ here), is the only biochemical reaction observed under the physiological conditions normally encountered by the Gram-negative bacterium *E. coli.* Under standard conditions (pH 7, 298 K, 1 bar pressure and 1 mol.L^−1^ substrate/product concentrations) the standard redox potential (*E*^0′^) of CO_2_/formate has been calculated as −420 mV, which is very close to H+/H_2_ where *E*^0′^ −410 mV (Reeve *et al*., 2017). The bacterium only produces this enzyme when environmental conditions favour the forward reaction – that is relatively high formate concentration and pH below 7 (Bohm *et al*., 1990). The volatility of the H_2_ product, and the rapid reactivity of the dissolved CO_2_ product to form carbonic acid and bicarbonate, keep the product concentrations low, which helps to increase the thermodynamic favourability of the forward reaction. Indeed, increasing the temperature that this reaction is performed at so increases the theoretical free energy available and allows some hyperthermophilic microbes to couple FHL activity to the generation of a transmembrane ion gradient that will support life (Lim et al., 2014).

Previous studies suggested that the FHL complex could potentially perform the ‘reverse’ reaction’ given the behaviour of the individual enzyme components of FHL (Sawers et al., 1985, Maeda et al., 2007, Bassegoda et al., 2014, McDowall et al., 2014) and early work in intact cells (Woods, 1936). Indeed, it has been suggested that an evolutionary progenitor of FHL – perhaps itself driven by a transmembrane ion gradient, or perhaps already under permissive conditions in the deep ocean – could be responsible for hydrogen-dependent CO_2_ fixation on early Earth (Nitschke & Russell, 2009). In this work, it was considered that the close standard redox potentials of the two half reactions of FHL, and evidence that the enzyme activity was not coupled to other biochemical processes such as electrochemical gradients (Pinske & Sargent, 2016), should allow the correct environmental conditions to be found that would drive the reverse reaction: *i.e.* increased pH; increased gas pressure; increased substrate concentrations; and rapid removal of the product from the vicinity of the enzyme.

Initial modelling was performed to allow prediction of the increases in aqueous concentrations of 1:1 mixtures of H_2_ and CO_2_ that could be obtained under applied pressure. This led to the design and testing of a novel bioreactor system that would allow sampling of cells under constant pressure and controlled pH. When headspace gas pressure was applied to a washed suspension of *E. coli* cells already containing FHL, the efficiency of the hydrogen-dependent CO_2_ reduction reaction was found to peak around 100 %. Indeed, in some cases calculations suggested slightly more formate was produced than CO_2_ gas was consumed (Figure 6). The HPLC-based method to quantify formate is consistent and reliable (Figure 4), therefore one likely explanation is that there is slight experiment-to-experiment variation in the substrate gas composition and associated pressure measurements, or that alternative sources of CO_2_ are present in the cells. Indeed, it should be considered that the biomass used here is extensively washed and placed in anaerobic buffer with no carbon or energy sources. The cells are effectively starving and it is possible breakdown of endogenous lipids or amino acids will generate some CO_2_ that could itself be reduced to formate.

The predicted concentrations of dissolved CO_2_ and H_2_ in solution in a 1:1 CO_2_:H_2_ headspace gas mixture at 10 bar pressure is 122.88 mmol.L^−1^ CO_2_ and 3.61 mmol.L^−1^ H_2_. Precise quantification of the gas mixture in the “ballast vessel” suggested instead that a 56:44 CO_2_:H_2_ mixture was present, representing 137.62 mmol.L^−1^ CO_2_ in solution. The *K*_m_ for CO_2_ for the formate dehydrogenase component of FHL is not known, however its *K*_m_ for formate is 26 mmol.L^−1^ (Sawers, 1994) and the reverse reaction has been studied by electrochemistry using 10 mmol.L^−1^ carbonate as an alternative substrate (Bassegoda et al., 2014). The *K*_m_ for H_2_ of the Hyd-3 [NiFe]-hydrogenase component has been estimated by electrochemistry techniques as 34 μmol.L^−1^ at pH 6 (McDowall et al., 2014). Thus, it can be concluded that at least the dissolved levels of the H_2_ substrate are clearly saturating under these test conditions. Note also that proton reduction activity by Hyd-3 is affected by direct product inhibition, with an inhibition constant calculated at 1.48 mmol.L^−1^ H_2_ (McDowall et al., 2014). This means Hyd-3 is likely to be biased towards H_2_ oxidation under the high pressure reaction conditions used here.

### Formate production and excretion from the cell

The accumulation of >500 mmol.L^−1^ (>0.5 Molar) formate in solution is quite remarkable (Figure 5a). This value far exceeds the substrate concentrations used. It should be noted well, however, that the formic acid accumulates outside of the cells in this experiment (Figure 1b). This immediate excretion of the formate from the cell upon its generation would be predicted to maintain product concentrations at low levels in the cell cytoplasm and so help maintain the hydrogen-dependent CO_2_ reduction activity. The most likely route for formic acid excretion is *via* the FocA channel (Waight *et al*., 2010, Wang *et al*., 2009). The mechanism of FocA is not yet fully agreed upon, with some hypotheses supporting a pH-gating mechanism where import is favoured at pH <7 and export is favoured, or perhaps FocA operates as a passive channel, at pH >7 (Lv *et al*., 2013, Lu *et al*., 2013, Lu *et al*., 2011, Lu *et al*., 2012). Recent work suggests FocA may function as an obligate formic acid/proton symporter at pH < 7 and therefore formate uptake into the cell may be driven by the protonmotive force (Wiechert & Beitz, 2017). In the key experiment described here (**Figure 5a**), the external environment is maintained at pH 8. If FocA is considered an open passive channel at alkaline pH (Lu et al., 2011), then the formic acid (p*K*_a_ = 3.75) produced in the cell cytoplasm, which is normally maintained at a pH 7.2-7.8 (Padan *et al*., 2005), will be drawn to and accumulate in the alkaline extracellular environment at a 10-times higher concentration than that in the cytoplasm for every pH unit difference (Nicholls & Ferguson, 2013).

### The possibility of using genetic and biochemical engineering to improve reaction rates

In this work, the rates and total amounts of formate produced from CO_2_ and H_2_ were compared in two *E. coli* strains with different formate metabolic capabilities (Figure 7). In addition to FHL, the *E. coli* FTD89 strain contains two quinone-dependent respiratory Fdhs named FDH-N and FDH-O (Sawers, 1994), which would be expected to consume periplasmic formate if a suitable electron acceptor was present. The *fdhE* mutation included in the RT1 strain would block assembly of these two enzymes (Schlindwein & Mandrand, 1991). The FTD89 strain also contains an active PFL enzyme. PFL is responsible for endogenous formate production from pyruvate under anaerobic conditions with the concomitant generation of acetyl CoA (Sawers & Watson, 1998). In the presence of a source of acetyl CoA, for example acetate and ATP, PFL can operate in reverse and generate pyruvate from formate (Zelcbuch et al., 2016). The *pflA* mutation included in the RT1 strain should negate all PFL activity in the *E. coli* cell (Crain & Broderick, 2014).

Although all efforts were made to produce washed cell suspensions free of oxygen and other chemical contamination, at small scale and using 1 bar gaseous substrates, removal of additional formate metabolism pathways seems to benefit both the yield of formate and its rate of production (Figure 7). Moreover, when the two strains were assayed in the high pressure system, the RT1 strain produced ~0.15 mmol formate. mg^−1^ total cellular protein (25 g cells at 10 bar) as a final concentration (Figure 8a), while the FTD89 strain that retains the other pathways produced ~0.12 mmol. formate. mg^−1^ total cellular protein (25 g cells at 10 bar) as a final concentration (Figure 6). The difference is small but may be important if maximum formate yield is the main aim of the carbon dioxide reduction process.

Additional genetic modifications may also improve the rate of CO_2_ reduction to formate. Although it was possible to optimise the amount of biomass used in the current experimental set-up, the current experiments are restricted to the native levels of enzyme that can be generated in the cell. One alternative approach would be to design a dedicated bacterial cell factory, maximally loaded with active FHL. All of the factors required for the biosynthesis of FHL are well understood (Sargent, 2016), and *E. coli* presents the ideal host organism for genetic engineering of strong promoters upstream of the required genes. One paradox is that the *E. coli* FHL enzyme must be membrane bound in order to be enzymatically active (Pinske & Sargent, 2016), which may limit total cellular levels of any overproduced enzyme unless mutations affecting membrane biosynthesis are also incorporated into any cell factory strain (Miroux & Walker, 1996).

Biochemical approaches may also be useful in optimising the rate of hydrogen-dependent CO_2_ reduction. The native formate dehydrogenase component of FHL contains a molybdenum atom at its active site, however it should be considered that tungsten-containing formate dehydrogenases may be better suited to CO_2_ reduction (Maia *et al*., 2015). Successful substitution of molybdenum by tungsten could reduce the midpoint potential of the metal centre significantly (Hagedoorn *et al*., 2003) that would, in theory, simultaneously drive electron flow towards CO_2_ while imposing a thermodynamic barrier on the opposite formate oxidation reaction from taking place.

## Conclusions

In summary, this study reports the use of high-pressure reactors for effective and efficient whole-cell biocatalysis by *E. coli.* The system could be considered a carbon capture technology, since the original aim was to process gaseous CO_2_ into a manageable product. Alternatively, the system may be considered as a specific formate generation technology. Taken altogether, our results demonstrate the production of >0.5 mol.L^−1^ formate with a rate exceeding 1 g of formate produced *per* litre *per* hour using only CO_2_ and H_2_ at 10 bar pressure. Moreover, bioconversion of CO_2_ with 100 % efficiency can be achieved. This approach does not require a large amount of biomass for effective conversion and the use of a well-known industrial workhorse organism such as *E. coli* presents several advantages for the production of such whole-cell biocatalysts and the opportunity to integrate this system into other bioprocessing projects.

The work provides proof-of-concept that FHL could be harnessed as a straightforward carbon capture device or CO_2_ recycling technology for industry. For direct use in heavy industry, however, the presence and impact of contaminant waste gases, such as carbon monoxide, should be considered. CO is a classic competitive inhibitor of [NiFe]-hydrogenases, but *E. coli* Hyd-3 has been observed to exhibit greater tolerance to CO attack than other enzymes, especially under H_2_ oxidation conditions (McDowall et al., 2014). This natural property, together with the potential to engineer heterologous enzymes that will metabolise any CO present (Gregg *et al*., 2016), means that the presence of CO in off gases is a problem that could be solved.

*E. coli* FHL could be employed as a means to specifically generate formate, which is a commodity in itself, can be directly used as H_2_ carrier or energy store (Jens *et al*., 2016), or can serve as feedstock for a wide range (bio)chemical reactions (Yishai *et al*., 2016). Alternatively, the formate so-produced could possibly be further converted to other products by incorporating recombinant enzymes into host organisms, representing a promising solution that couples the recycling of CO_2_ to its use as carbon source and chemical feedstock (Aresta & Dibenedetto, 2007). The experiments described here have been conducted on non-growing cell suspensions. Genetic engineering has recently demonstrated the ability of modified *E. coli* to grow on exogenous formate as a carbon source (Zelcbuch et al., 2016, Yishai *et al.*, 2017). This raises the possibility that FHL activity, as a source of formate from gaseous CO_2_, could be incorporated into growing cells to allow CO_2_ assimilation into biomass and other bio-products.

Finally, hydrogen gas is the obligate source of reductant in this biological system. For future applications, the use of renewable hydrogen sources would be ideal, however in these laboratory-based experiments H_2_ produced by steam reformation of natural gas has been used. This is not sustainable in the long term, however it should be considered that for every CH_4_ chemically processed one CO_2_ and four H_2_ are generated. In terms of the FHL enzyme, a single H_2_ is required to reduce CO_2_ to formate. Thus, by employing the pressurised *E. coli* FHL system described here it could be possible to render such chemical generation of H_2_carbon-neutral by capturing the CO_2_ waste yet retain production of excess H_2_ for use in other applications.

## EXPERIMENTAL PROCEDURES

### Bacterial strains and growth conditions

The *E. coli* K-12 strains used in this study (Table 1) were FTD89 (Δ*hyaB*, Δ*hybC*) (Sargent *et al*., 1999), RT1 (Δ*hyaB*, Δ*hybC*, Δ*pflA*, Δ*fdhE*) (Pinske & Sargent, 2016) and the FHL-minus control strain RT2 (Δ*hyaB*, Δ*hybC*, Δ*pflA*, Δ*fdhE*, Δ*hycA*-*I*::Kan^R^) (Pinske & Sargent, 2016). Anaerobic fermentative growth was performed in sealed bottles at 37 ^<u>°</u>^C for 12-14 hours using TYEP medium (Begg *et al*., 1977), pH 6.5, containing 0.8% (w/v) glucose and 0.2% (w/v) sodium formate to induce the expression of the FHL regulon. In the case of the *E. coli* RT2 strain, kanamycin was added in the culture medium at a final concentration of 50 μg.mL^−1^.

**Table 1:** *E. coli* K-12 strains used in this study.

| Strain | Relevant Genotype | Reference |
| --- | --- | --- |
| MC4100 | *E. coli* K-12: F<sup>-</sup>, [*araD139*]B/r, $\Delta(\textit{argF-lac})169^*$, $\lambda^-$, e14<sup>-</sup>, *flhD5301*, $\Delta(\textit{fruK-yeiR})725$ (*fruA25*), *relA1*, *rpsL150*(str<sup>R</sup>), *rbsR22*, $\Delta(\textit{fimB-fimE})632(::\text{IS1})$, *deoC1* | (Casadaban & Cohen, 1979, Peters *et al.*, 2003) |
| FTD89 | as MC4100, $\Delta\textit{hyaB}$, $\Delta\textit{hybC}$ | (Sargent *et al.*, 1999) |
| RT1 | as MC4100, $\Delta\textit{hyaB}$, $\Delta\textit{hybC}$, $\Delta\textit{pflA}$, $\Delta\textit{fdhE}$ | (Pinske & Sargent, 2016) |
| RT2 | as MC4100, $\Delta\textit{hyaB}$, $\Delta\textit{hybC}$, $\Delta\textit{pflA}$, $\Delta\textit{fdhE}$, $\Delta\textit{hycA-l}::\text{Kan}^R$ | (Pinske & Sargent, 2016) |

### Small scale catalysis of hydrogen-dependent CO_2_ reduction to formate at ambient pressure

After anaerobic fermentative growth, 1 L of culture was harvested by centrifugation (Beckman J6-MI centrifuge) for 30 min at 5000 g and 4 ^<u>°</u>^C. The cell paste was then washed twice by suspending the cell pellet in 20 mmol.L^−1^ 3-(N-morpholino)propanesulfonic acid (MOPS) buffer pH 7.4 and centrifuging (Fisher Scientific Accuspin 1R centrifuge) for 15 minutes at 3000 *g* and 4 ^<u>°</u>^C. Finally, the cell pellet was suspended in the same buffer at 50 μ.L^−1^ (wet weight). Next, 500 μL of the washed whole-cell suspension, corresponding to 25 mg of wet cells, was transferred to a Hungate tube containing 2.5 mL of MOPS buffer. The tubes were sealed and flushed with argon for 5 min, then flushed with H_2_ for 5 min before 5 mL CO_2_ was added to the tubes. The cells were incubated at 37 ^<u>°</u>^C for 23 hours. Samples of the clarified liquid phase were analysed by HPLC.

### Larger scale experimental setup for the high-pressure reactor

The experiments were carried out in two identical, stainless steel 1.2 L volume Premex reactors used as a ‘production vessel’ and gas mixture ‘ballast vessel’ (see Fig. S2, ESI^‡^). The reactors are fitted with customised gas-entraining mechanical stirrers, temperature and pressure probes, internal cooling coils (mainswater) and fluidised jacket (connected to a Huber 405w thermostatic bath), the latter ensuring that isothermal conditions between production and ballast vessels can be maintained. The temperature and pressure was continuously monitored, controlled and data logged by a Procontrol Ordino process interface. High pressure pH and reference probes (Corr Instruments Inc.) were added to the production vessel and pH changes were monitored over the time course of the reaction using the Rosemount 56 Emerson advanced analyser. The ballast vessel was connected to the bioreactor *via* a stainless steel transfer line equipped with a back pressure regulator to ensure constant pressure gas feed. Feeding of base (sodium hydroxide 1.0–2.0 M) was conducted *via* a Knauer HPLC-pump K-120 connected to the production vessel with Ar back pressure. The pump rate was set up at 2.5 mL.min^−1^ at the beginning of the experiment and then controlled manually in order to maintain the pH above 6.8. At the start of an experiment both vessels were heated to 110 ^<u>°</u>^C under vacuum for 2 hr, cooled to 37 ^<u>°</u>^C (operational conditions) and back-filled with Ar to ensure removal of oxygen and moisture. The vessels were purged with Ar another 3-times by filling to 10 bar before being vented (< 1 bar pressure). The H_2_:CO_2_ gas ballast vessel was prepared by pressurising the reactor with first CO_2_ and then H_2_ at 40 bar total pressure maintaining the fixed pressure ratio of ca. 1:1 at 37 ^<u>°</u>^C and 500 rpm. The gas composition was confirmed by Agilent GC-TCD (thermal conductivity detector).

The production vessel was prepared as follows. After anaerobic fermentative growth, cultures were harvested by centrifugation (Beckman J6-MI centrifuge) for 30 min at 5000 g and 4 ^<u>°</u>^C. The cell extract was then washed twice by suspending the cell pellet in either 20 mmol.L^−1^ MOPS pH 7.4 or 200 mmol.L^−1^ Tris.HCl pH 8.0 and centrifuged (Fisher Scientific Accuspin 1R centrifuge) for 15 min at 3000 g and 4 ^<u>°</u>^C. The cell pellet was weighed before being suspended in the same buffer at a final amount of 50 g.L^−1^ (wet weight), unless otherwise stated in the figure legends. Next, 500 mL of washed whole-cells was transferred into the production vessel and purged with argon for 30 min at 37 ^<u>°</u>^C and 500 rpm. Finally, the reaction was initiated by pressurising the transfer line and the production vessel with the H_2_:CO_2_ mixture at 2, 4, 6 or 10 bar pressure. The production vessel pressure was maintained constant over the time course of the reaction (~23 hours) at the desired pressure by a back pressure regulator connected to the transfer line. Samples of the liquid phase in the production vessel are collected at different time points, filtered (0.2 μm PES filters) and analysed without further dilution by HPLC (equipped with UV and RI detectors).

### Substrate calculations

For the small scale experiments conducted at ambient pressure, the substrate calculations were made as in Pinske *et al.* (2016). For the larger scale experiments conducted using high-pressure reactors, the concentration of gases in the liquid phase was calculated by considering Henry’s law using gas constants at 298K/25C to be 1282.1 L.atm.mol^−1^ and 29.4 L.atm.mol^−1^ for H_2_ and CO_2_, respectively, and calculating values at 310K/37C using the equation:

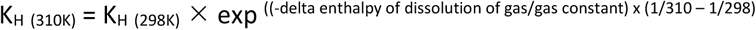

giving derived gas constants at 310K/37C of 1373.4 L.atm.mol^−1^ and 39.9 L.atm.mol^−1^ for H_2_ and CO_2_, respectively. A H_2_:CO_2_ gas mixture of composition (44:56 ratio determined experimentally in this work) at 2, 4, 6 or 10 bar pressure corresponds to 27.52, 55.05, 82.58 and 137.63 mmol.L^−1^ CO_2_ in the aqueous phase, respectively. Note that under the stirring rates employed (~500 rpm) it cannot be assumed that 100 % of gas applied moves to the liquid phase. The efficiency of CO_2_ conversion was calculated by determining the moles of CO_2_ consumed in the H_2_:CO_2_ ballast vessel during the reaction and comparing to the amount of formic acid produced. The moles of CO_2_ consumed were determined according to the ideal gas law considering (i) the H_2_:CO_2_ mixture is an ideal gas with a compressibility factor (Z) of 1.00000288; (ii) the gas mixture is composed of ~44 % H_2_ and ~56 % CO_2_ as determined by TCD analysis.

### Analytical methods

Total cell protein amount was estimated based on the OD_600_ of the culture and the assumption that 1 L culture with an OD_600_ of 1 contains 0.25 g of dry cell of which half is assumed to be protein. Organic acid analysis and quantification was determined by HPLC using either a Dionex UltiMate 3000 system equipped with an Aminex HPX-87H column (BioRad) or a Shimadzu Prominence HPLC equipped with a Rezex ROA-Organic Acid H^+^ (8%) LC Column 300 × 7.8 mm and Synergi 4 μm Hydro-RP 80Å, LC-column 150 × 4.6 mm (Phenomenex). Samples of 10 or 100 μL that were previously clarified through 0.2 μm filters were applied to the columns equilibrated in 5 mmol.L^−1^ H_2_SO_4_ with a flow of 0.5 mL.min^−1^ at either 50 ^<u>°</u>^C/30 min/UV (210 nm) detection (Dionex system) or 40 ^<u>°</u>^C/30 min/RI detection (Shimadzu system). The formate eluted at either 16.2 min or 19.5 min, respectively. The composition of the gas mixture was confirmed by Agilent GC-TCD (thermal conductivity detector) and a standard curve of formic acid (1-500 mmol.L^−1^) was prepared.

## ACKNOWLEDGEMENTS

We thank the Biotechnology and Biological Sciences Research Council (BBSRC) C1net Network in Industrial Biotechnology & Bioenergy (NIBB) for proof-of-concept funding (award POC-8-sargent-C1net).

